# SENIES: DNA Shape Enhanced Two-layer Deep Learning Predictor for the Identification of Enhancers and Their Strength

**DOI:** 10.1101/2021.05.14.444093

**Authors:** Ye Li, Fanhui Kong, Hui Cui, Chunquan Li, Jiquan Ma

## Abstract

The identification of enhancers has always been an important task in bioinformatics owing to their major role in regulating gene expression. For this reason, many computational algorithms devoted to enhancer identification have been put forward over the years. To boost the performance of their methods, more features are extracted from the single DNA sequences and integrated to develop an ensemble classifier. Nevertheless, the sequence-derived features used in previous studies can hardly provide the 3D structure information of DNA sequences, which is regarded as an important factor affecting the binding preferences of transcription factors to regulatory elements like enhancers. Given that, we here propose SENIES, a DNA shape enhanced deep learning predictor, for the identification of enhancers and their strength. The predictor consists of two layers where the first layer is for enhancer and non-enhancer identification, and the second layer is for predicting the strength of enhancers. Besides utilizing two common sequence-derived features (i.e. one-hot and *k*-mer) as input, it introduces DNA shape for describing the 3D structures of DNA sequences. Performance comparison with state-of-the-art methods conducted on the same datasets demonstrates the effectiveness and robustness of our method. The code implementation of our predictor is publicly available at https://github.com/hlju-liye/SENIES.

## I. INTRODUCTION

**T**HE living and development of organisms are inseparable from the proper function of gene expression in cells, which is regulated by the concerted cooperation of various types of gene regulatory elements located in the non-coding regions of genome [1]. The typical regulatory elements include enhancers, promoters, silencers, insulators and so on. Among these, enhancers are deemed as the most crucial ones responsible for regulating the transcription of their target genes. Different from gene proximal regulatory elements like promoters, the location of enhancers relative to their target genes cannot be simply formulated. They can be located upstream, downstream, or sometimes within introns of their target genes. Beyond that, some enhancers can bypass their nearest genes to regulate more distant ones along a chromosome. In some special cases, they can even regulate genes on another chromosome [2]. This kind of locational uncertainty has made their prediction a challenging task for modern biologists.

The early attempts to identify enhancers on a genome-wide scale started with biologically experimental methods. The most representative one is chromatin immunoprecipitation followed by deep sequencing (ChIP-seq) [3]. By targeting at enhancer associated marks like transcription activator p300 [4], histone H3 monomethylated at K4 (H3K4me1) [5] and H3 acetylated at lysine 27 (H3K27ac) [6], this technique has been successfully applied in some cell lines to identify enhancers. Another category of the experimental method is based on chromatin accessibility, i.e., detecting the open regions on the genome. Two commonly used techniques are DNase I digestion coupled to sequencing (DNase-seq) [7] and transposase-accessible chromatin followed by sequencing (ATAC-seq) [8]. Further studies on enhancers have shown that they can function as transcriptional units and produce non-coding RNAs (eRNAs), which are hallmarks of active enhancers [9]. However, eRNAs are generally unstable and have a short half-life, making them extremely hard to detect in cells. Despite this, several techniques have been developed to detect the expression of eRNAs, such as global run-on sequencing (GRO-seq) [10] and cap-analysis of gene expression (CAGE) [11]. While all the aforementioned experimental methods are useful to some extent, there is currently no golden standard in biology for enhancer identification. And beyond that, experimental ways are time-consuming, labor-intensive, and impractical at present to be generally applied to all cell types at various stages.

On this account, some computational methods have been put forward to fill this gap. As the first attempt, Heintzman et al. [5] developed a computational prediction algorithm to locate enhancers in the ENCODE regions of HeLa cells based on similarity to chromatin profiles in the training set. Firpi et al. [12] further proposed a computational framework named CSI-ANN. It was composed of a data transformation and a feature extraction step followed by a classification step with time-delay neural network. With the discovery and map of increasing amount of histone modifications, the selection of the optimal set from the entire range of chromatin modifications for enhancer identification becomes an urgent task. So Rajagopals et al. [13] developed RFECS, a Random-Forest based algorithm, to integrate 24 histone modification profiles in all for the identification of enhancers in several cell types. They claimed that their method not only led to more accurate predictions but also identified the most informative and robust set of three chromatin marks for enhancer identification. However, the common goal of the aforementioned methods was simply to label a DNA sequence as an enhancer or not. They neglected to determine the strength of enhancers, i.e., their activity level, which is also biologically meaningful. Given that, Liu et al. [14] proposed a two-layer predictor named iEnhancer-2L by formulating DNA elements with pseudo *k*-tuple nucleotide composition. In the second layer of their predictor, they identified enhancers’ strength for the first time. Considering the unsatisfied performance of iEnhancer-2L, they further improved their algorithm by formulating sequences with different feature representations and using ensemble learning in their later study iEnhancer-EL [15]. Based on Liu et al.’s work, Jia et al. [16] developed a predictor called EnhancerPred by extracting three types of sequence-based features and using support vector machine (SVM) to identify enhancers and their strength. Recently, Cai et al. [17] proposed a more advanced predictor named iEnhancer-XG where five different feature representations derived from sequences were used as the input of XG-boost, a new learning algorithm based on gradient boosted decision trees.

Over the past few years, deep learning has seen a comprehensive penetration into various fields, including computer vision, natural language processing and even bioinformatics [18–23]. Naturally, a torrent of deep learning based methods for enhancer identification have sprung up, such as EP-DNN [24], BiRen [25], DECRES [26], PEDLA [27], DeepEnhancer [28]. Comparing to traditional machine learning methods, deep learning obviates the needs for manually curating features and can unearth informative hidden patterns in the data. While these deep learning based enhancer predictors give much better performance than that of traditional machine learning based ones, their weaknesses are also quite obvious, i.e., the prediction of enhancer strength is not reflected in their methodologies.

We here present a two-layer enhancer predictor named SENIES for identifying not only enhancers (first layer), but also their strength (second layer). The predictor utilizes DNA shape information besides two common sequence-derived features (i.e. one-hot encoding and *k*-mer) as the input of our developed deep learning model. DNA shape refers to the three-dimensional (3D) structures of DNA. While several former studies have pointed out its role in the recognition of 3D DNA structures for transcription factors [29–31], the application of this particular feature has been limited to the modelling of TF-DNA binding [32–33]. In the process of gene regulation, the chromatin structures of enhancer regions tend to be open so as to provide a protein-binding platform for a combination of transcription factors and co-factors. Inspired by this, we hypothesize that DNA shape can be used as a complementary feature for enhancer identification. In the study of iEnhancer-2L, Liu et al. introduced six DNA local structural parameters [14]. However, these local structural parameters are just predicted from the physicochemical properties of two neighboring base pairs, which cannot accurately reflect the 3D structure of DNA sequences. Given this, we here use DNAshape [34], a Monte Carlo (MC) simulation based method to derive more informative DNA shape features. Comparing to existing state-of-the-art methods, the performance of SENIES gets an obvious boost for both layers of enhancer identification. More importantly, it proves that DNA shape can be used as another major feature to enhance the identification of enhancers and their strength.

## II. Materials AND Methods

### A. Benchmark and independent datasets

The benchmark and independent datasets used in our study were obtained from a series of previous works [14–17]. The two datasets were constructed based on chromatin state annotation. Specifically, Ernst *et al*. [35] mapped nine chromatin marks across nine cell types. They used recurrent combinations of these marks to define 15 chromatin states such as repressed, poised and active promoters, strong and weak enhancers. Then ChromHMM [36] was developed using a multivariate Hidden Markov Model to learn these chromatin states information and obtain the genome-wide chromatin state annotation. Afterwards, different sorts of DNA sequences were selected as candidate samples to construct the two datasets based on the genome-wide chromatin state annotation given by ChromHMM in these nine cell types.

A detailed description of the post-processing on these candidate samples can be looked up in the original works of Liu *et al* [14–15]. In this paper, we just report the composition of the final datasets. The benchmark dataset consists of 2968 sequence samples, where each of them is 200 bp long. They are used to construct the first layer predictor. Among these, 1484 samples are enhancers, and the remaining 1484 samples are non-enhancers. Of the 1484 enhancer samples, strong enhancers and weak enhancers are half and half. They are used to construct the second layer predictor. The independent dataset was constructed using the same protocol as the benchmark dataset. It is composed of 200 enhancers and 200 non-enhancers. Half of the 200 enhancers are strong enhancers, and the other half are weak enhancers. The independent dataset is used to evaluate the performance of predictors developed using the benchmark dataset. It is noted that there is no overlap between the samples in the benchmark dataset and the independent dataset.

### B. Feature representation

A fundamental problem in developing a bioinformatics related predictor is how to formulate a biological sequence (i.e. DNA sequence in our study) with some specific feature representations. In truth, quite a lot of feature representation methods for biological sequences have been proposed so far, such as *k*-mer, pseudo *k*-tuple nucleotide composition (PseKNC), subsequence and mismatch profile [37]. In this study, we explore three feature representation methods, namely, one-hot, *k*-mer and DNA shape. Among these, one-hot and *k*-mer represent two common ways to formulate DNA sequences, which have been widely used in bioinformatics to resolve various types of classification and prediction problems. However, sequence-constraint methods often fail to identify non-coding functional elements like enhancers because they neglect to consider the 3D structures of DNA sequences [38]. Hence, DNA shape is used as a supplemental feature for providing 3D structural information. A detailed description of these three different feature representation methods is as below.

#### 1) K-mer

Oligomers of length *k*, or *k*-mers, refer to all the subsequences of length *k* contained within a biological sequence [39]. It is a widely used and probably the simplest feature representation method for modeling the properties and functions of biological sequences [37]. In the case of deoxyribonucleic acid, each sequence comprises four different types of nucleotides (i.e. A, C, G and T). *K*-mer approach first lists all the possible subsequences of length *k* and scan the whole DNA sequence to find the occurrence frequency of each subsequence. Then the occurrence frequency of each subsequence is combined by the order of the listed subsequences. This combined feature vector is called the *k*-mer feature vector of that sequence. Given a sequence ***S***, then the *k*-mer feature vector of ***S*** can be defined as:

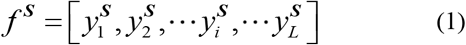

where 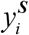 is the occurrence frequency of the *i*th *k* neighboring nucleotides in the sequence ***S***, and the value of *L* is 4^*k*^. As pointed in [40], selecting the parameter *k* in *k*-mer feature representation is difficult due to its inherent limitation. The *k*-mer feature vector tends to become sparser and encode less effective information when *k* gradually increases. We observed that the *k*-mer vector is quite sparse when the value of *k* is greater than 5 in our case since the length of enhancer sequences in our dataset is relatively short. Thus, we use a combination of different *k*-mer feature vectors where *k* ranges from 1 to 5 and concatenate their results as a final feature vector.

#### 2) One-hot encoding

One-hot encoding is another common feature representation method for formulating DNA sequences. This encoding scheme is especially popular when it is combined in use with convolutional neural networks (CNN) [20–23, 28]. With this feature representation, every nucleotide in the sequence will be transformed into a four-dimensional binary vector, where the bit marking the current nucleotide is set to one, and all the other bits are set to zero. By such, four nucleotides, A (adenine), C (cytosine), G (guanine), and T (thymine) will be transformed into binary vectors (1, 0, 0, 0) ^⊤^, (0, 1, 0, 0) ^⊤^, (0, 0, 1, 0) ^⊤^, and (0, 0, 0, 1) ^⊤^ respectively. Then the binary vector for each nucleotide in the DNA sequences will be merged into a binary matrix. Given a DNA sequence ***S =*** *N*_1_ *N*_2_ ⋯ *N*_*i*_ ⋯ *N*_*L*_, then the one-hot encoding matrix ***M*** for sequence ***S*** can be formulated as:

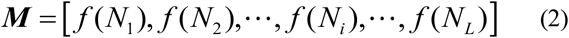

where *f* is a function defined as:

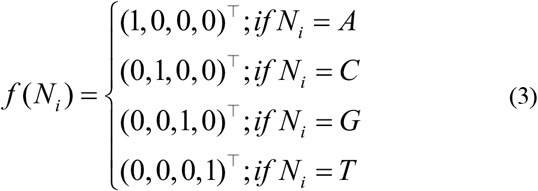

#### 3) DNA shape

DNA shape presents the chromatin structural information of DNA sequences, which the above two feature representation methods cannot provide. Zhou *et al*. [34] proposed a high-throughput method called DNAshape for predicting chromatin structural features from DNA sequences. The core of their methodology is a pentamer based model built from all-atom Monte Carlo simulations where a sliding-window approach is used to mine DNA shape features from DNA sequences. Later, Chiu *et al*. [41] developed DNAshapeR, an R/Bioconductor package based on DNAshape to generate DNA shape predictions in an ultra-fast, high-throughput and user-friendly manner. At first, only four DNA shape features were included, that is minor groove width (MGW), helix twist (HelT), propeller twist (ProT) and Roll, for their extremely importance in the recognition of DNA structures. In their latest package release, another 9 DNA shape features were added, and the entire repertoire was finally expanded to a total of 13. Among these, seven features were nucleotide shape parameters and the other six were base pair-step parameters. An illustration of the distinction between generating the two types of DNA shape parameters is presented in Fig. 1 where the DNA sequence is scanned with a pentamer sliding window. For each pentamer subsequence currently being scanned, a DNA shape prediction value of the central nucleotide or two prediction values of the two central base pair steps will be computed based on the specific type of the given DNA shape parameter. Supposing the length of a DNA sequence is N, then the dimension of nucleotide shape parameter-based feature vector and base pair-step shape parameter-based feature vector can be easily induced as N−4 and N−3, respectively. Since there are 7 nucleotide shape parameters and 6 base pair-step shape parameters used in our study, the length of the concatenated shape feature vector can be formulated as 7×(N−4) + 6×(N−3). Given N=200 in our case, an input DNA sequence can be finally encoded with a DNA shape vector of 2554 dimensions.

**Fig. 1.**
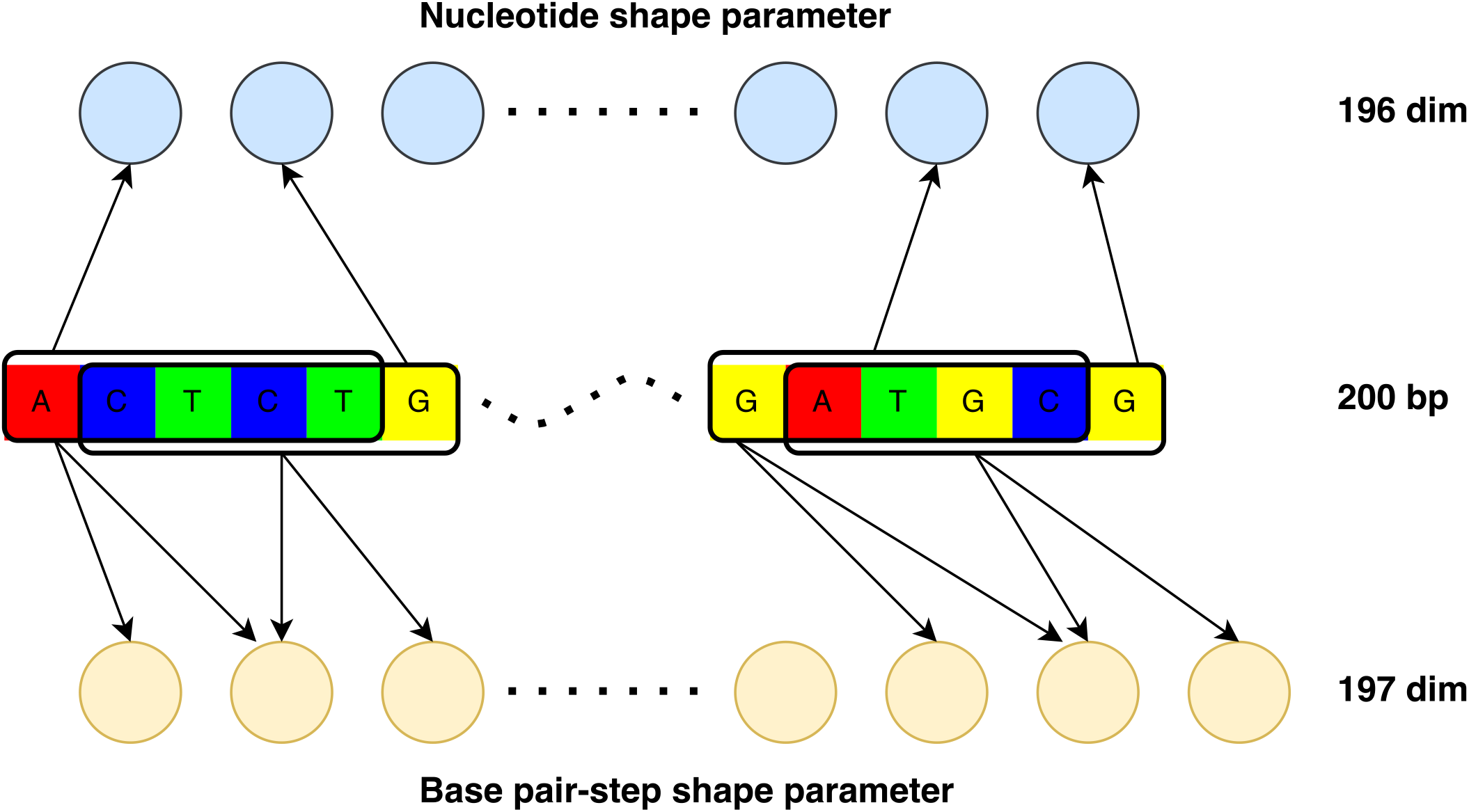
A simple sketch on illustrating the difference between the generation of nucleotide shape parameters and base pair-step shape parameters. The DNA sequence is scanned with a pentamer sliding window to derive DNA shape feature vectors. For each pentamer subsequence being scanned, a single prediction value of the central nucleotide will be computed for nucleotide parameters like MGW and ProT. While for base pair-step shape parameters like Roll and HelT, the prediction values of the two central base pair steps will be provided. Taking the first pentamer in the sequence as an example, the central nucleotide is T and the two central base pair steps are CT and TC respectively. It’s worth noting that the second central base pair step of a pentamer subsequence is identical to the first central base pair step of the next pentamer subsequence, so they share the same DNA shape prediction value.

### C. Network architecture

The network architecture of our designed deep learning model is mainly composed of four modules (Fig. 2). The top three modules, from left, are DNA shape module, one-hot module and *k*-mer module, respectively. In DNA shape module, the R package DNAshapeR is used to generate the shape feature predictions from the given set of DNA shape parameters as the protocol illustrated in Fig. 1. In our study, all the 13 DNA shape features were generated for each DNA sequence. In one-hot encoding module, the input sequence is firstly encoded with a 4×200 one-hot binary matrix and then fed into a two-layer convolution neural network (CNN). The first layer consists of 16 convolutional kernels of size 4×8, and the second layer is composed of 32 convolutional kernels of size 1×8. Notably, while it has not been shown in the figure for simplicity, each convolutional layer is followed by a max-pooling layer with dimension 1×2. After convolutions, the output feature maps are flattened and concatenated to a feature vector. In *k*-mer module, the frequency vectors of different *k*-mers (*k* ranges from 1 to 5) are computed separately and then concatenated. Finally, output vectors of the top three modules are concatenated and sent into the joint module based on a multilayer perceptron network (MLP) with two hidden layers. The number of neurons in the first and second layers is 512 and 64, respectively. The output layer has only one neuron representing the binary classification result of our model.

**Fig. 2.**
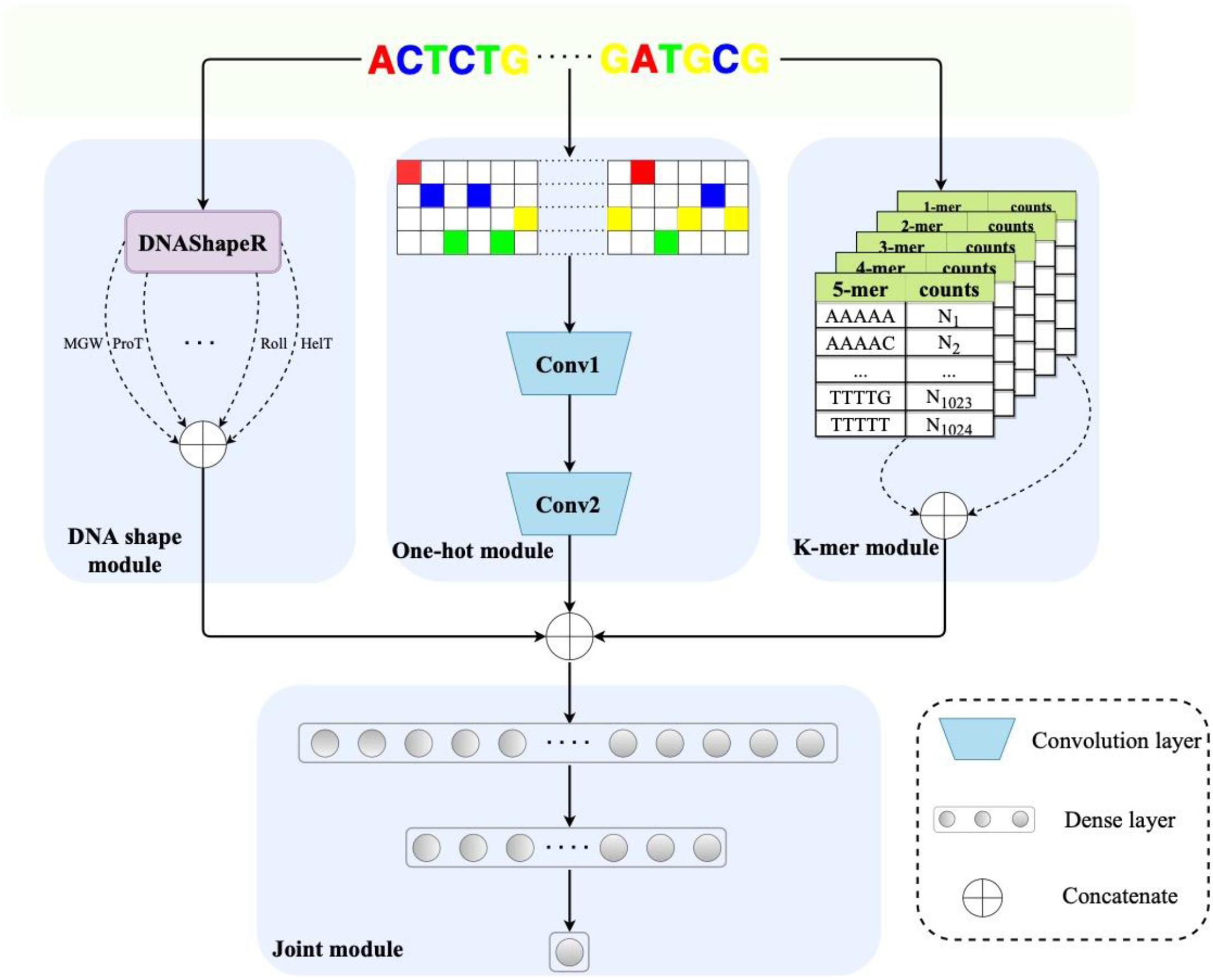
The network architecture of our designed deep learning model. The top three modules, from left, are DNA shape module, one-hot module and *k*-mer module respectively and the bottom one is the joint module. The output vector from the top three modules will be concatenated and fed into the joint module where a multilayer perceptron (MLP) is used to get the final prediction result.

### D. Implementation details

All the models built in our study are implemented with Pytorch 1.4.0 [42]. Adam [43] is used as the optimizer, and the learning rate for training models is set to 1e-5. The mini-batch size is 20. Binary cross entropy is employed as the loss function. To prevent models from overfitting, an early stopping strategy is used with patience set to 30, and the evaluation metric MCC is monitored on validation set. That is to say, after a successive of 30 training epochs with no increase on the metric Matthew correlation coefficient (MCC), the training process is stopped, and the model with the highest MCC value is saved for further evaluation on the independent dataset.

### E. Performance evaluation

For a fair comparison, the identical set of evaluation metrics as used in a series of previous works is also adopted in our study.

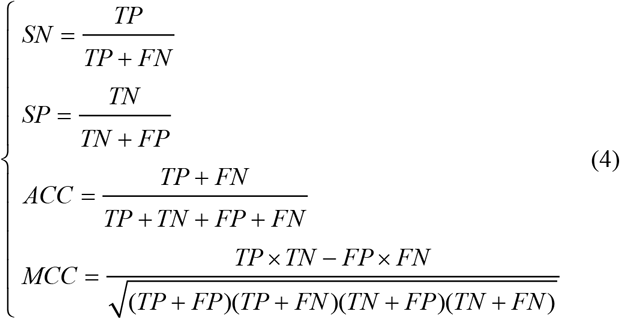

These metrics are sensitivity (SN), specificity (SP), accuracy (ACC), and MCC. They are all formulated in (4) where TP, TN, FP, and FN represent the number of true positives, true negatives, false positives and false negatives, respectively.

Considering that our dataset is relatively small, a five-fold cross validation is used to evaluate the performance of our model. Specifically, our benchmark dataset is partitioned into five subsets. In each fold, four parts of them are used as the training set, and the remaining one is used for validation. This process is repeated after five times and each time we will get a different data partition for the training of our model. Then the five trained models will be tested in turn on the independent dataset. Since there are five predictions from five-folds, we adopt an ensemble learning strategy to get the final prediction on the independent dataset (i.e. taking the mean value of the five prediction results). To validate the stability of our models, the five-fold cross-validation experiments are conducted ten times with all the results shown with box plots.

## III. Results AND Discussion

### A. Investigation of basic and full sets of DNA shape features

DNAshape initially provided the prediction of only 4 DNA shape features (MGW, ProT, Roll and HelT), which we refer to as basic set. Another 9 features (Rise, Roll, Shift, Slide, Tilt, Buckle, Opening, Shear, Stagger and Stretch) were added in the latest version of DNAshapeR and the feature set was expanded to a total of 13, which we refer to as full set. While former studies focused mainly on the basic set, the full set of DNA shape features were used in our study. Naturally, it is worth evaluating whether these 9 new added DNA shape features have a positive effect on the identification of enhancers and their strength. To prove it, we compared the performance of basic set and full set of DNA shape features on the independent dataset. For the first layer targeting at distinguishing enhancers from non-enhancers, the performance of full set is better than that of basic set by a narrow margin on four evaluation metrics except for SN (Fig. 3a). For the second layer aiming to distinguish strong enhancers from weak enhancers, the edge of full set comparing to basic set is more obvious with a lead of performance on all five evaluation metrics (Fig. 3b). Thus, the introduction of the additional 9 DNA shape features has a positive influence for both layers of enhancer prediction, especially the second layer.

**Fig. 3.**
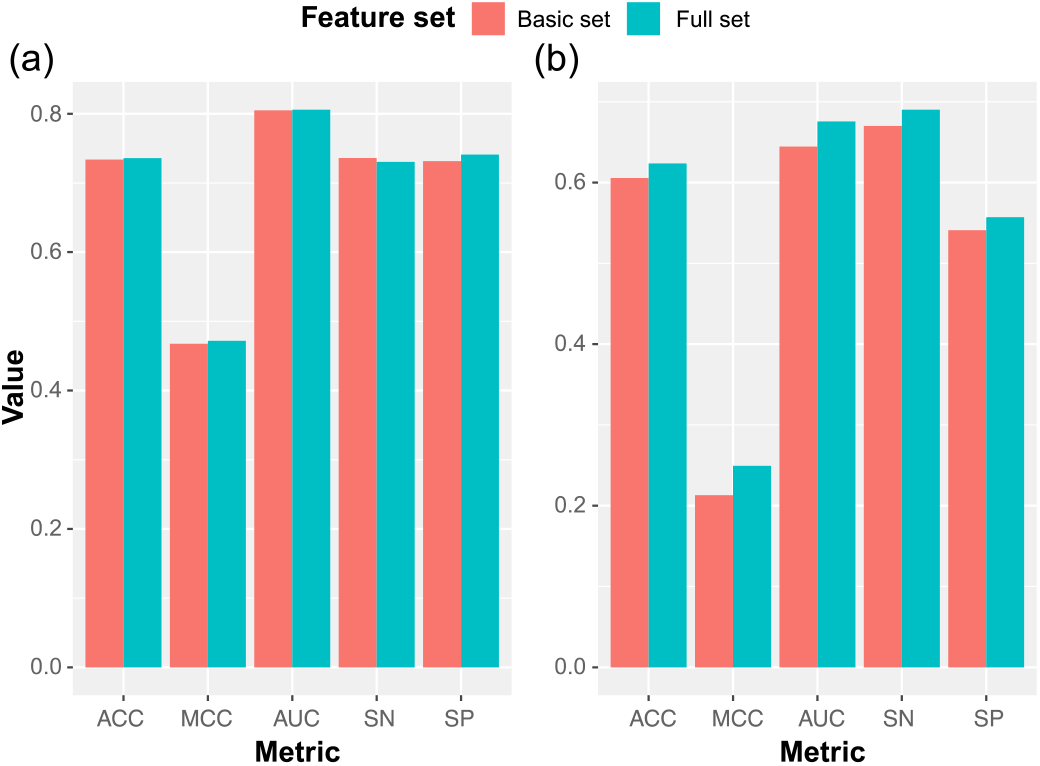
Performance comparison between the basic set and full set of DNA shape features on the (**A**) first layer and the (**B**) second layer.

### B. Visualization of DNA shape features

We use aggregated line plots to visualize DNA shape features of the positive and negative samples. The line plots of some representative shape features have been chosen and shown for the first and second layer (Figs. 4, 5). For MGW, the two aggregated lines are separated on the first layer while overlapped on the second layer. This finding suggests that MGW effectively differentiates enhancers from non-enhancers but may not be ideal for distinguishing strong enhancers from weak enhancers. For the Opening shape feature, the two aggregated lines are separated on both layers suggesting its effectiveness for both layers of enhancer identification. For Buckle, though the aggregated line of the positive set is higher than that of the negative set, there is overlap between them. And for Tilt, the profiles of both aggregated lines fluctuate around zero, suggesting this shape feature can be hardly used for enhancer identification task. The aggregated line plots of the other DNA shape features are attached as supplemental files.

**Fig. 4.**
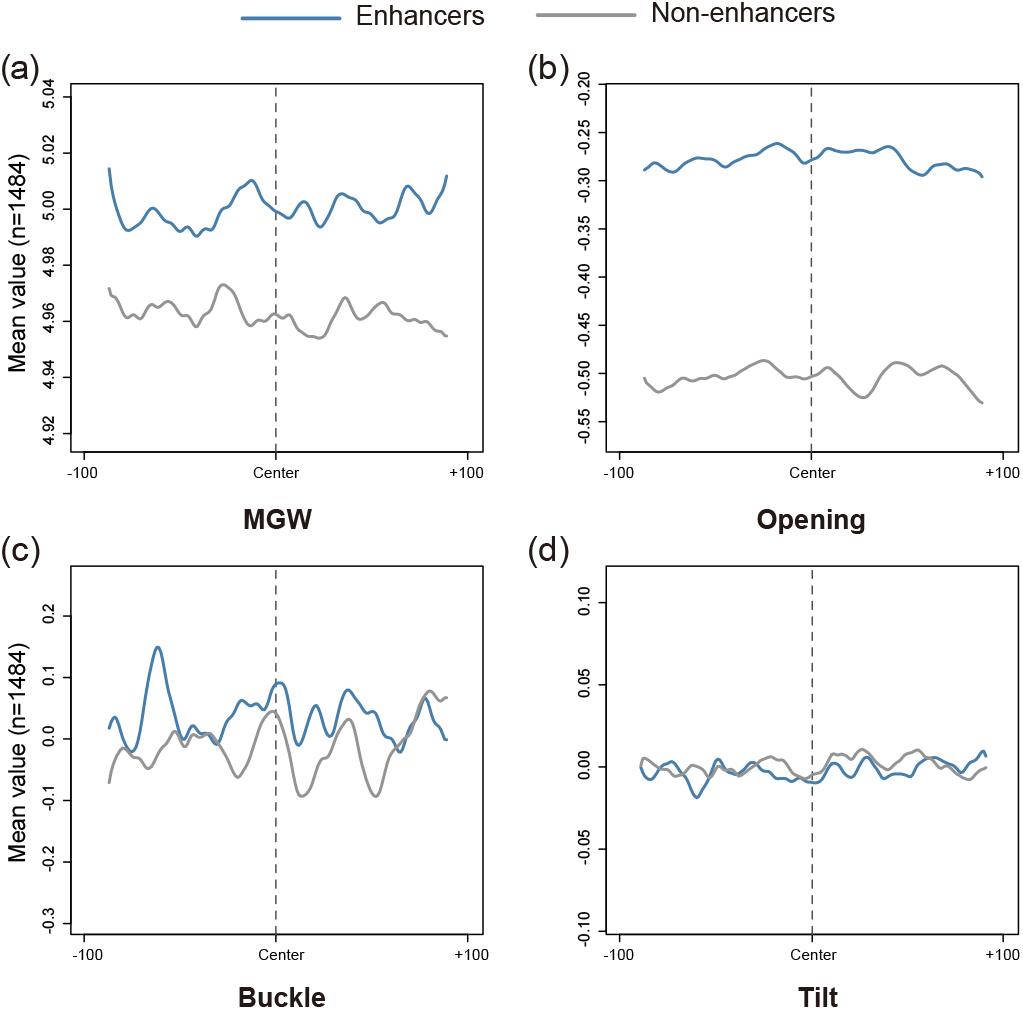
Visualization of four representative DNA shape feature features (**A**) MGW (**B**) Opening (**C**) Buckle (**D**) Tilt with aggregated line plots on the first layer.

**Fig. 5.**
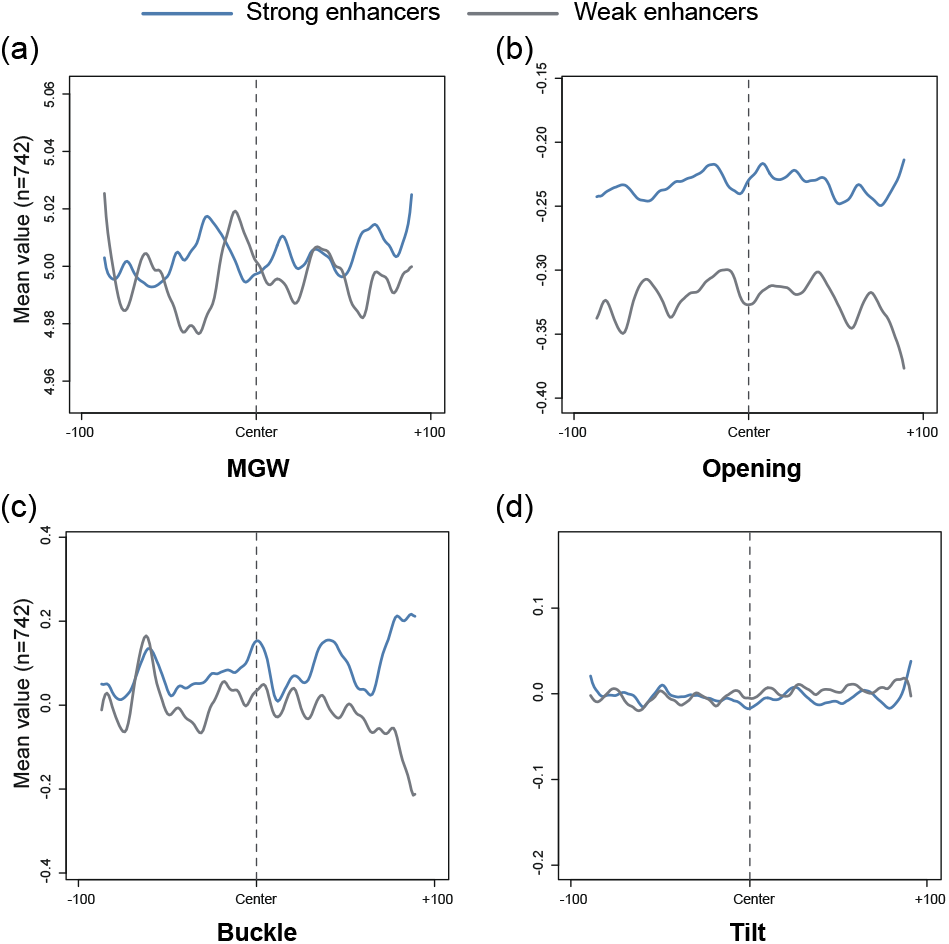
Visualization of four representative DNA shape feature features (**A**) MGW (**B**) Opening (**C**) Buckle (**D**) Tilt with aggregated line plots on the second layer.

### C. Performance of different feature representations and their combinations

As there are total three different types of feature representations used in our method, a natural question in face of us is to determine the importance of them in the identification of enhancers. For that, each module used in Fig. 2 is taken out and designed as an individual network. Besides, different modules are combined to see the effect of various combinations of these feature representations. For the sake of simplicity, the model combining one-hot and DNA shape modules is denoted by ‘O-S’; the model combining one-hot and *k*-mer modules is denoted by ‘O-K’; the model combining DNA shape and *k*-mer modules is denoted by ‘S-K’; the model combining all three modules is denoted by ‘ALL’. Then the performance of all these models is evaluated with a five-fold cross validation on the independent dataset. To reflect the stability of these models, the cross-validation experiment is repeated after 10 times. The results of the ten experiments are shown with box plots (Fig. 6). The average results are given in Table I and Table II.

**Table I.** Performance of different feature representations and their combinations on the independent dataset of the first layer.

| Feature representations | ACC (%) | MCC | AUC | SN (%) | SP (%) |
| --- | --- | --- | --- | --- | --- |
| One-hot | 73.70 | 0.475 | 0.812 | 71.20 | 76.20 |
| Shape | 73.55 | 0.471 | 0.806 | 74.00 | 73.10 |
| K-mer | 75.80 | 0.516 | 0.830 | 76.00 | 75.60 |
| O-S | 74.13 | 0.483 | 0.814 | 72.85 | 75.40 |
| O-K | 76.15 | 0.523 | 0.834 | 76.25 | 76.05 |
| S-K | 76.40 | 0.528 | 0.833 | **77.25** | 75.55 |
| All | **76.80** | **0.536** | **0.834** | 76.90 | **76.70** |

The highest value achieved on every single metric has already been marked in bold.

**Table II.** Performance of different feature representations and their combinations on the independent dataset of the second layer.

| Feature representations | ACC (%) | MCC | AUC | SN (%) | SP (%) |
| --- | --- | --- | --- | --- | --- |
| One-hot | 57.95 | 0.163 | 0.638 | 66.40 | 49.50 |
| Shape | 62.05 | 0.243 | 0.679 | 68.40 | **55.70** |
| K-mer | 62.50 | 0.256 | 0.681 | 73.30 | 51.80 |
| O-S | 63.05 | 0.276 | 0.661 | 78.90 | 47.20 |
| O-K | 65.45 | 0.316 | 0.712 | 75.50 | 55.40 |
| S-K | 67.35 | 0.358 | 0.739 | 79.20 | 55.50 |
| All | **68.25** | **0.380** | **0.753** | **82.00** | 54.50 |

The highest value achieved on every single metric has already been marked in bold.

**Fig. 6.**
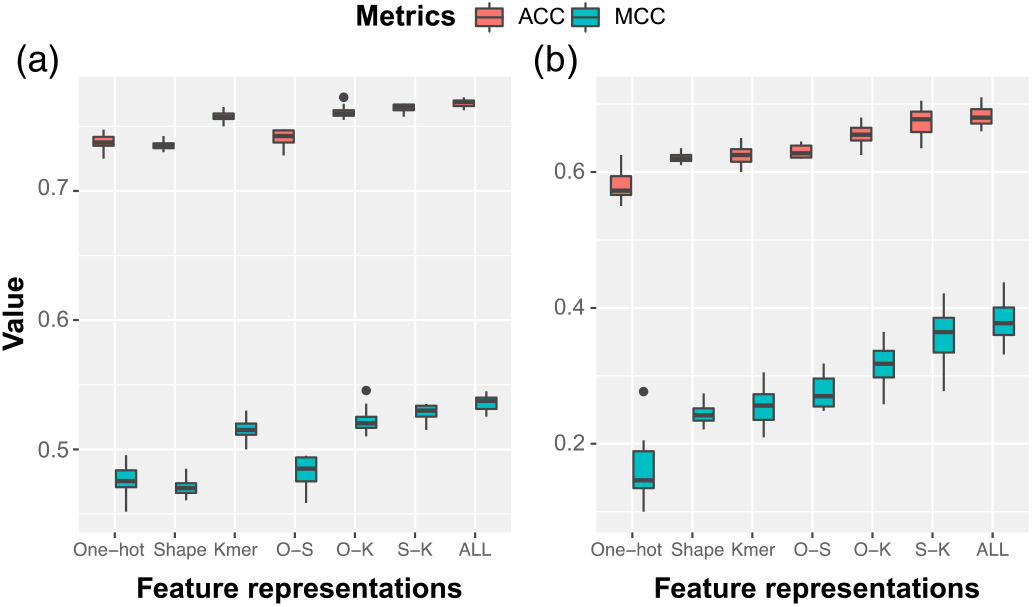
Performance of different feature representations and their combinations.

As shown, for the first layer, *k*-mer approach achieves the best performance among the three single feature representations with ACC of 75.80% and MCC of 0.516. The performances of the other two feature representations, one-hot and DNA shape, are very close, where one-hot obtains an ACC of 73.70% and MCC of 0.475, and DNA shape obtains an ACC of 73.55% and MCC of 0.471. For the second layer prediction, the performance of one-hot feature is unsatisfactory, with ACC of 57.95% and MCC of 0.163. For the other two feature representations, the performance of DNA shape is slightly lower than that of *k*-mer. The ACC and MCC of them are 62.05%, 62.50% and 0.243, 0.256, respectively. Overall, *k*-mer approach outperforms the other two feature representation methods on both layers, followed by DNA shape and one-hot.

As for those combined feature representations, their performance basically all improved compared to that of single feature representations. Notably, the model combining all three feature representations performed best on both layers. For the first layer, it achieves the best performance on all evaluation metrics except for SN. And for the second layer, it outperforms the other models on all evaluation metrics except for SP. For that reason, it is selected as our final model for performance comparison with state-of-the-art methods.

### D. Performance comparison with existing methods

We compare the performance of our final model on the independent dataset with some existing state-of-the-art methods (Table III). As shown, our method achieves the best performance on most evaluation metrics for both layers. Note that ACC and MCC are deemed as the two most important ones among the five evaluation metrics for our prediction task [15]. For the first layer, ACC is improved by 1.4% and MCC is improved by 4.1%. And for the second layer, these two metrics are boosted by 7.5% and 39.7%. As another important evaluation metric for binary classification, AUC is improved from 0.817 to 0.834 for the first, and from 0.680 to 0.753 for the second layer, respectively. Overall, the performance of our method surpasses these existing methods terms of a comprehensive comparison on these evaluation metrics.

**Table III.** Performance comparison with existing methods on the independent dataset.

| Task | Method | ACC (%) | MCC | AUC | SN (%) | SP (%) |
| --- | --- | --- | --- | --- | --- | --- |
| 1st layer | iEnhancer-2L | 73.00 | 0.460 | 0.806 | 71.00 | 75.00 |
|  | Enhancer-Pred | 74.00 | 0.480 | 0.801 | 73.50 | 74.50 |
|  | iEnhancer-EL | 74.75 | 0.496 | 0.817 | 71.00 | **78.50** |
|  | iEnhancer-XG | 75.75 | 0.515 | - | 74.00 | 77.50 |
|  | SENIES | **76.80** | **0.536** | **0.834** | **76.90** | 76.70 |
| 2nd layer | iEnhancer-2L | 60.50 | 0.218 | 0.668 | 47.00 | **74.00** |
|  | Enhancer-Pred | 55.00 | 0.102 | 0.579 | 45.00 | 65.00 |
|  | iEnhancer-EL | 61.00 | 0.222 | 0.680 | 54.00 | 68.00 |
|  | iEnhancer-XG | 63.50 | 0.272 | - | 70.00 | 57.00 |
|  | SENIES | **68.25** | **0.380** | **0.753** | **82.00** | 54.50 |

The highest value achieved on every single metric has already been marked in bold.

## IV. Conclusion

As the core parts of DNA regulatory elements responsible for regulating gene expression, enhancers have always been paid the most attention in bioinformatics. However, their locational uncertainty and the poor understanding of their sequence code have made their identification an extremely challenging task [2]. To further enhance the identification of enhancers and their strength, most of the current machine learning based predictors tend to derive more feature representations from DNA sequences and ensemble the predictions of individual features. While some of the existing structural biology and genomics studies have confirmed the relationship between TF-DNA binding and the recognition of chromatin structures, we observe that the sequence-derived feature representations used in previous works cannot reflect the structural information of DNA sequences. In light of this, DNA shape is used as an additional feature input besides two commonly used ones, i.e., one-hot and k-mer. Through the ablation experiments, we find that the performance of feature combined models is boosted comparing to those single feature representation based ones. Above all, the deep learning model with all the three feature representations as input achieves the best performance for both layers of enhancer identification. And in the comparison with some state-of-the-art methods, SENIES achieves remarkable performance with only three features all derived from DNA sequences. Yet, our study can be further improved by integrating epigenetic features like histone modifications. Many studies have associated the enhancer activities with certain characteristic histone modification patterns [5, 13, 44]. However, histone marks are cell line specific while the enhancer dataset used in our study is not, so it is beyond our reach at present. Nevertheless, it does not affect us to apply this method in later constructed cell line specific enhancer dataset.

## Acknowledgment

We thank Hui Cui for checking and improving the English language in this paper. We also thank Bin Liu for making the enhancer dataset public.

---

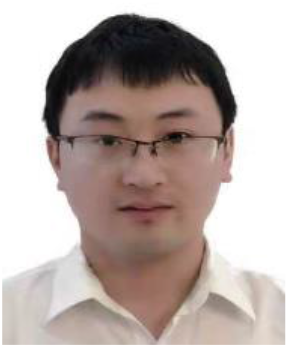

**Ye Li** received his bachelor degree (2017) in information management and information system from China Pharmaceutical University. He is now pursuing the M.S degree in computer science and technology at Heilongjiang university, China. His research interests include bioinformatics and deep learning.

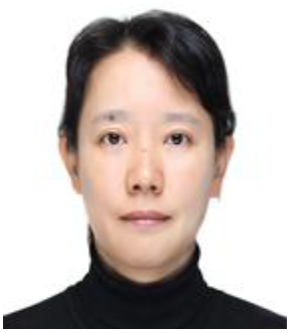

**Fanhui Kong** received her Ph.D (2016) in instrument science and technology from Harbin Institute of Technology in China. She is currently associate professor of department of data science and technology in Heilongjiang University. Her research interests are in medical image analysis and deep learning.

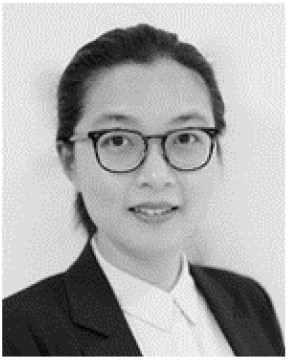

**Hui Cui** received PhD and MPhil degrees in computer science from The University of Sydney, Australia, in 2016 and 2013, and BEng degree in electrical engineering from Harbin Institute of Technology, China, in 2011. She is currently a Lecturer at the Department of Comp Sci & Info Tech, La Trobe University, Australia. She was a research fellow at the Biomedical & Multimedia Information Technology (BMIT) Research Group, The University of Sydney (2016-2018). Her research area is in the interdisciplinary field of machine Learning, image processing, medical image analysis, and health informatics.

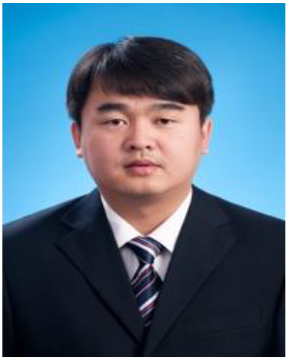

**Chunquan Li** received his Ph.D (2012) in Biophysics from Harbin Medical University in China. He is currently professor and deputy chair of School of Medical Informatics in Harbin Medical University, Daqing campus. His research subject is the construction and analysis of biomolecular network and pathway model for complex diseases. His research interests are in bioinformatics and system biology.

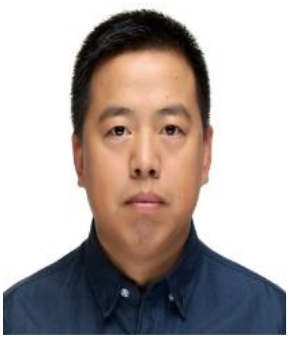

**Jiquan Ma** received his Ph.D (2010) in computer science from Harbin Institute of Technology in China. He was a post doctoral research scholar at Harbin Medical University in China (2012-2015). He is currently chair of department of computer science and technology in Heilongjiang University. He was a visit scholar at the University of North Carolina at Chapel Hill, USA (2018-2019). His research interests are in computer vision, medical image analysis and deep learning.

